# *In vivo* Imaging β-cell Function Reveals Two Waves of β-cell Maturation

**DOI:** 10.1101/159673

**Authors:** Jia Zhao, Weijian Zong, Yi Wu, Jiayu Shen, Dongzhou Gou, Yiwen Zhao, Runlong Wu, Fuzeng Niu, Xu Wang, Xuan Zheng, Aimin Wang, Yunfeng Zhang, Jing-Wei Xiong, Liangyi Chen, Yanmei Liu

## Abstract

The insulin-secreting cells generated from stem cells *in vitro* are less glucose responsive than primary β-cells. To search for the missing ingredients that are needed for β-cell maturation, we have longitudinally monitored function of every β-cell in Tg (*ins:Rcamp1.07*) zebrafish embryos with a newly-invented two-photon light-sheet microscope. We have shown that β-cell maturation begins from the islet mantle and propagates to the islet core during the hatching period, coordinated by the islet vascularization. Lower concentration of glucose is optimal to initiate β-cell maturation, while increased glucose delivery to every cell through microcirculation is required for functional boosting of the β-cells. Both the initiation and the boosting of β-cell maturation demands activation of calcineurin/NFAT by glucose. Calcineurin activator combined with glucose promotes mouse neonatal β-cells cultured *in vitro* to mature to a functional state similar to adult β-cells, suggesting a new strategy for improving stem cell-derived β-like cell function *in vitro*.

## Introduction

Pancreatic β-cells secrete insulin to regulate blood glucose metabolism. An insufficient functional β-cell mass causes glucose intolerance and diabetes. To generate new therapeutic approaches for diabetes, β-cell development has been intensively studied during the last two decades. While much has been understood regarding the early development of pancreatic progenitor cells (Pan & Wright, 2011), the mechanisms regarding the final β-cell maturation are still poorly defined (Kushner et al, 2014). β-cells reside in islets that consist of endocrine, vascular, neuronal and mesenchymal cells. Various cues arising from this neurovascular milieu, such as gap junctions, neuronal transmitters, endothelial factors and hormones, have been reported to be involved in β-cell development (Borden et al, 2013; Carvalho et al, 2010; Cleaver & Dor, 2012; Omar et al, 2016). The absence of these factors from the culture media that is used by current *in vitro* differentiation protocols may account for the limited glucose-responsiveness of stem cell-derived β-like cells (Bruin et al, 2015; Rezania et al, 2014). Hence, knowing how pancreatic β-cells mature *in situ* during development may guide the addition of key factors to the culture medium to promote the final maturation of β-like cells *in vitro.*

A major difficulty in studying β-cell maturity *in vivo* with traditional methods is to evaluate β-cell function independent of β-cell mass. Imaging function of individual β-cells will overcome this problem. The primary function of a mature β-cell is to quickly secrete the stored insulin in responding to increases of blood glucose concentration. Glucose-triggered Ca^2+^ influx in pancreatic β-cells is essential for the insulin secretion, thus often serves as a functional marker to evaluate β-cell maturity in living organisms (Pagliuca et al, 2014; Rezania et al, 2014). Ca^2+^ transient in primary β-cells can be imaged in isolated islets when β-cells are labelled with genetically encoded fluorescent Ca^2+^ indicators. However, to image Ca^2+^ transients in β-cells *in vivo* noninvasively remains the unmet challenge due to the non-transparency of the pancreas tissues in mammals.

Here we used zebrafish as the model animal in which β-cell development is conserved with mammals and their maturation processes can be observed in the transparent, externally developed embryos of zebrafish (Huang et al, 2001). With every β-cell labelled with Rcamp1.07, a red fluorescent Ca^2+^ indicator in Tg (*ins:Rcamp1.07*) zebrafish, we use our newly developed high-resolution two-photon three-axis digital scanning light-sheet microscope (2P3A-DSLM) to visualize glucose-stimulated Ca^2+^ responses in individual β-cells *in vivo*. The first glucose-responsive β-cells appear from the islet mantle at 48 hours postfertilization (hpf). Based on the Ca^2+^ transient kinetics, we characterized the glucose-responsive β-cells into two stages: the initiation and the boosting of the maturation. This early β-cell maturation of the islet mantle requires low local glucose concentration and is independent of vascularization. With islet vascularization from 60 hpf to 72 hpf, increased glucose concentration delivered through islet microcirculation, initiates β-cell maturation in the islet core and pushes all glucose-responsive β-cells towards further maturity. By manipulating with inhibitors, activators and the dominant negative mutant, we demonstrate that the calcineurin/NFAT signaling pathway acts downstream of glucose to initiate, boost and sustain β-cells maturation. Finally, we showed that neonatal mouse β-cells within isolated islets can be promoted towards optimal maturity *in vitro* by the combination of a calcineurin activator and glucose, demonstrating that mammalian β-cell maturation shares the conserved mechanism. This data also highlights the practicality of direct activation of calcineurin/NFAT in pushing stem cell-derived β-like cells towards optimal maturity *in vitro*.

## Results

### Visualization of β-cell maturation in vivo with 2P3A-DSLM

To visualize β-cell function, we created a transgenic zebrafish line, Tg (*ins:Rcamp1.07),* in which the red fluorescent calcium indicator Rcamp1.07 (Ohkura et al, 2012) is under the control of the insulin promoter. Rcamp1.07 is expressed exclusively in β-cells, as confirmed by the cellular co-localization of Rcamp1.07 with EGFP in Tg (*ins:Rcamp1.07);* Tg (*ins:EGFP*) double transgenic fish (Huang et al, 2001) (Figure 1-Figure Supplement 1A-F) and with immunofluorescently labelled insulin in Tg (*ins:Rcamp1.07*) fish (Figure 1-Figure Supplement 1G-J). Within 5 min after stimulating live 72 hpf Tg (*ins:Rcamp1.07*) fish embryos with 20 mM glucose, we observed a robust transient increase in the fluorescence intensity of Rcamp1.07 under a wide-field microscope (Figure 1-Figure Supplement 2 and Supplemental Movie 1), indicative of the glucose-stimulated Ca^2+^ transients *in vivo*. However, individual β-cells are hardly discernible under the wide-field (Figure 1-Figure Supplement 2) or a single-photon selective-plane illuminative microscope (1P-SPIM) (Figure 1-Figure Supplement 2). With a two-photon microscope (TPM), we could resolve individual β-cells in the XY plane, but the cell boundaries along Z-axis were blurred due to the low axial resolution and high scattering of the illumination light in deep tissues. Only with our newly-developed 2P3A-DSLM (Supplemental Movie 2), we could reconstruct a clear three dimensional structure of the islet in live fish embryos (Figure 1A and B, Supplemental Movie 2 and Supplemental Movie 3). To our knowledge, this is the first time to achieve high-resolution *in vivo* imaging of individual β-cells and their functions.

**Figure 1.**
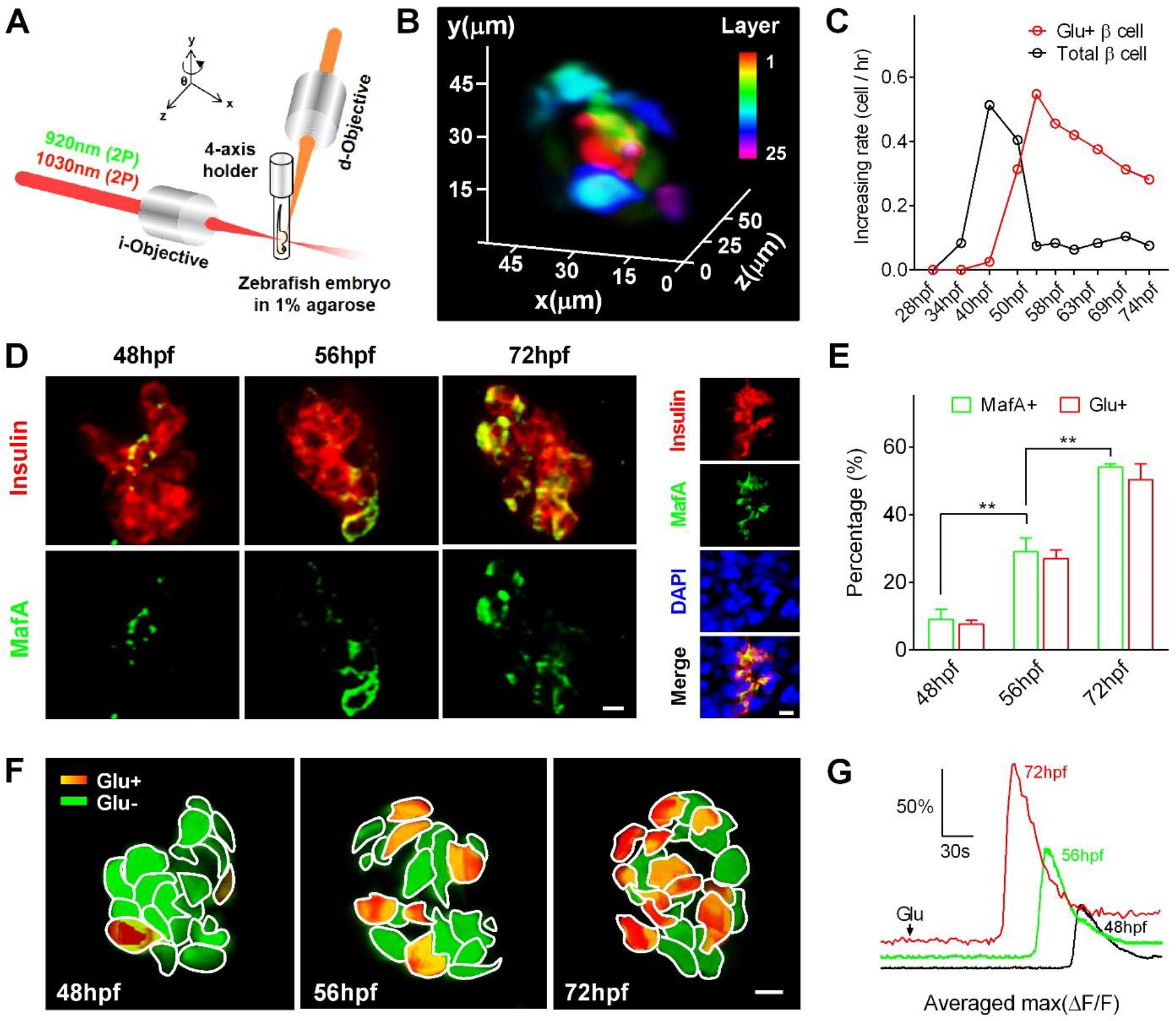
Visualization and characterization of the β-cell maturation process *in vivo* by 2P3A-DSLM. (*A*) An abbreviated scheme of the dual-colour 2P3A-DSLM imaging system. (*B*) 3D-projection of all pancreatic β-cells in a living 72 hpf *Tg(ins:Rcamp1.07*) zebrafish embryo. The different colours in the colour bar represent different depth of the islet. (*C*) The increasing rates of total β-cell and glucose-responsive β-cell numbers at different stages from 28 to 74 hpf. n = 10-16 embryos for each stage. (*D*) Whole-mount immunofluorescent labelling of insulin and MafA in embryos at 48, 56 and 72 hpf (left). Representative immunofluorescent labelling images of cryostat sections of 72 hpf zebrafish embryos (right). (E) Quantifications of MafA-positive β-cells and glucose-responsive β-cells at the indicated stages. n = 46 embryos for each condition. \*\**P*<0.01. (*F*) An illustration of glucose-responsive (red) and glucose nonresponsive (green) β-cells of the islets in live *Tg(ins:Rcamp1.07) Tg(ins:EGFP*) embryos at 48, 56 and 72 hpf. (*G*) Average traces of glucose-triggered maximum Ca^2+^ transients at the indicated stages. n = 10-16 embryos for each condition. (Scale bars: 10 μm; scale bars apply to *D* and *F*.) See also Figure 1-Figure Supplement 1, 2 and 3, and Supplemental Movies 1, 2 and 3.

By counting insulin-positive cells and glucose-responsive β-cells at different stages of Tg (*ins:Rcamp1.07);* Tg (*ins.EGFP*) fish embryos, we identified a proliferation of β-cells during 36-48 hpf, followed by a progressive increase in glucose-responsive β-cells during the hatching period (Figure 1C and Figure 1-Figure Supplement 3A). Along with the increase of glucose-responsive β-cells, mature β-cell marker MafA positive β-cells also increased (Figure 1D and E), indicating that by counting glucose-responsive β-cells we could evaluate the *bona fide* maturation process of β-cells. Because the speed (time to rise) and the maximal amplitude (Max ΔF/F) of the glucose-stimulated Ca^2+^ transients are key parameters of β-cell functionality (Bruin et al, 2015; Rezania et al, 2014), we used them as criteria to evaluate the maturity levels of individual β-cells *in vivo* (Figure 1-Figure Supplement 3B). The first glucose-responsive β-cell appeared at 48 hpf and we defined it to be at the initiation stage of maturation. Compared to glucose-responsive β-cells at 48 hpf, the cells at 56 hpf exhibited faster and larger Ca^2+^ responses evoked by glucose stimulation on average as many cells have entered the function boosting stage. These average responses were further accelerated and enhanced at 72 hpf, when β-cells responded within ∼ 90 s of glucose administration with a ∼150% of the maximal amplitude of the evoked Ca^2+^ transient (Figure 1 F and G and Figure 1-Figure Supplement 3C and D). Therefore, with our β-cell-function visualization zebrafish system, we observed a gradual maturation process of β-cells *in vivo* from 48 to 72 hpf.

### Sequential maturation of β-cells from the islet mantle to the core is coordinated with islet vascularization

Next, we investigated whether islet vascularization affect β-cell maturation *in vivo* by labelling both β-cells and vascular endothelial cells in Tg (*ins:EGFP);* Tg (*flkl:imCherry*) fish (Wang et al, 2013). We found that blood vessels initiated contacts with the islet mantle from 48 to 60 hpf and further penetrated into the inner layers of the islet from 60 to 72 hpf (Figure 2A and Supplemental Movie 4). Most of the glucose-responsive β-cells was proximal to blood vessels (Figure 2-Figure Supplement 1A), and their longitudinal increase was paralleled with the increase of β-cells adjacent to the vessels (Figure 2B and Figure 2-Figure Supplement 1B).

**Figure 2.**
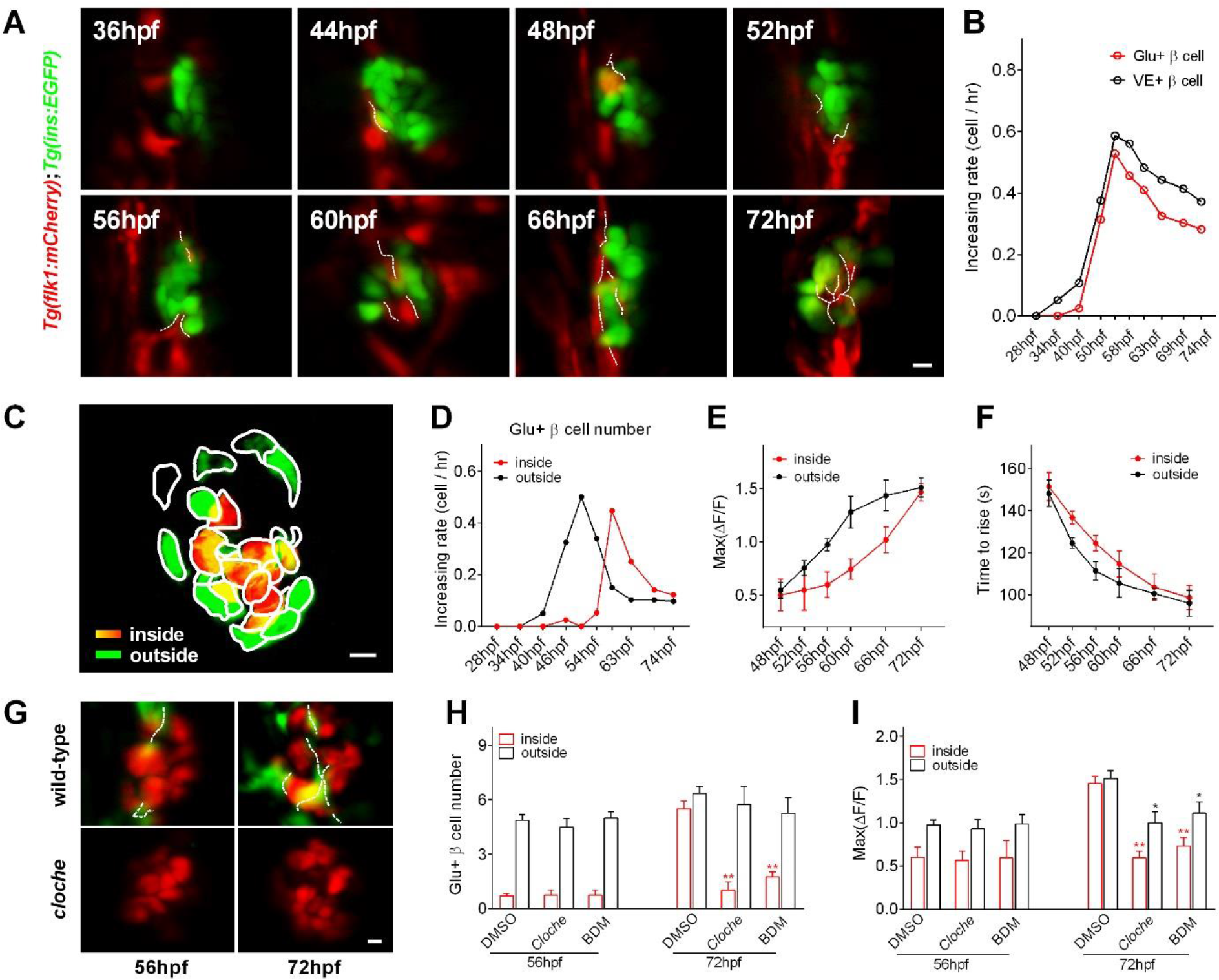
Sequential maturation of β-cells in the mantle and the core of the islet from the early to the late hatching period, which is coordinated with islet microcirculation. (*A*) Representative 3D-projections of β-cells (green) and neighbouring blood vessels (red) at different stages in live *Tg(ins:EGFP);Tg(flk1:mCherry*) embryos. (*B*) The comparison of the increasing rate of β-cells adjacent to the vascular endothelium (VE) with that of glucose-responsive β-cells at different stages from 28 to 74 hpf. n = 4-7 embryos for each stage. (*C*) An illumination of classification of β-cells into two populations, cells in the mantle (green) and in the core (red) of the islet. (*D*) Time-dependent increases in the glucose-responsive β-cells from the mantle (dark) and the core (red) of the islet. n = 10-16 embryos for each stage. (*E-F*) Time-dependent increases in the maximum amplitude (*E*) and the speed (*F*) of the glucose-triggered Ca^2+^ transients in β-cells from the mantle and the core of the islet. n = 10-16 embryos for each stage in *E* and *F*. (*G*) Representative 3D-projections of the β-cells (red) and neighbouring blood vessels (green) in live wild-type or *cloche^-/-^ Tg(ins:Rcamp1.07);Tg(flk1:GFP*) embryos. (*H-I*) Numbers of glucose-responsive β-cells (*H*) and their maximal Ca^2+^ responses to glucose (*I*) in the mantle and core of the islets in *cloche^-/-^* mutants and 2,3-BDM-treated embryos. n = 4-6 embryos for each stage in *H* and *I*. \**P*<0.05, \*\**P*<0.01. (Scale bars: 10 μm; scale bars apply to *A, C* and *G*.) See also Figure 2-Figure Supplement 1 and Supplemental Movie 4.

As islet vascularization progressed from the mantle to the core, we then examined whether β-cells in the mantle and β-cells in the core matured differently during development (Figure 2C). From 48 to 60 hpf, β-cells in the mantle acquired function earlier than those in the core, with more glucose-responsive β-cells (Figure 2D), higher maximum Ca^2+^ transients (Figure 2E) and faster responses (Figure 2F). Only during the late hatching period did β-cells in the core start to mature at an accelerated pace. At 72 hpf, they were indistinguishable from β-cells in the mantle in terms of all related parameters (Figure 2D-F). Therefore, β-cells in the mantle mature differently from those in the core both in the initiation time window and function boosting dynamics.

To explore whether the heterogeneous maturation process is caused by islet vascularization, we examined β-cell maturation in *Tg(ins:Rcamp1.07) cloche* mutant embryos, which have no vascular endothelial cells or blood cells but a normal number of β-cells (Figure 2G and Figure 2-Figure Supplement 1C) (Field et al, 2003a). Glucose-responsive β-cells in 56 hpf *cloche* embryos were indistinguishable from those in age-matched controls (Figure 2H and I). At 72 hpf, in contrast, *cloche* mutants had fewer glucose-responsive β-cells in the islet core and exhibited smaller maximum Ca^2+^ transients in responding β-cells than the controls (Figure 2H and I). Without islet microcirculation, β-cells of *cloche* mutants stayed in a maturation state similar to that in 60 hpf control embryos (Figure 3-Figure Supplement 1). Therefore, β-cells mature in two waves *in vivo:* first, mantle β-cells acquire glucose responsiveness independent of islet vascularization during 48-60 hpf; next, islet vascularization initiates the maturation of the β-cells in the islet core and promotes all glucose-responsive β-cells to further mature during 60-72 hpf. To test whether blood circulation or endothelial cells provide the key inductive factor, we stopped the blood circulation with 2,3- butanedione monoxime (2,3-BDM) (Bartman et al, 2004) for 9 hours in *ins:Rcamp1.07* embryos at either the early or the late hatching period. Despite intact islet vascularization after the circulation blockade (data not shown), 2,3-BDM treatment inhibited β-cell maturation to a similar extent as that of *cloche* mutants (Figure 2H and I). Therefore, blood circulation *per se,* but not endothelial cells, provides the key inductive signal for β-cells in the islet core to initiate and further promote their maturation.

### Increased glucose delivered by the microcirculation controls sequential β-cell maturation

Glucose, the major nutrient in the circulation system, is known to regulate embryonic pancreatic endocrine cell differentiation *in vitro* (Guillemain et al, 2007). To explore whether glucose could be the inductive signal for β-cell maturation, we used 3-mercaptopicolinic acid (3-MPA), an inhibitor of gluconeogenic phosphoenolpyruvate carboxykinase 1 (*pck1*), to suppress endogenous glucose synthesis (Jurczyk et al, 2011). This leads to severe and equal suppression of the maturation of β-cells in the mantle and the core of the islet during the whole hatching period (Figure 3 and Supplemental Movie 5). To probe the optimal endogenous glucose concentration required for β-cell maturation at the early and the late hatching periods, we incubated fish embryos with a combination of 3-MPA and different concentrations of exogenous glucose (3 mM, 5 mM, 8 mM or 20 mM). From 44 to 53 hpf, 3 mM glucose completely rescued the β-cell maturation defects caused by 3-MPA treatment, whereas 8 mM glucose only partially rescued the number of glucose-responsive β-cells in the islet mantle (Figure 3A and C). In contrast, from 60 to 69 hpf, we had to increase the exogenous glucose to 8 mM to fully rescue the phenotype (Figure 3A-D), and 3 mM glucose was insufficient to maintain a normal functional β-cell mass in the islet core (Figure 3A-D). During the entire hatching period, 20 mM glucose performed worst in the rescue experiments, indicative of glucotoxicity at this concentration, as reported in mammalian β-cells previously (Poitout et al, 2006). Therefore, different from vascularization, glucose is required for both β-cells in the islet core and those in the mantle for maturation initiation, function boosting and maintenance. On the other hand, our results also demonstrate that islet vascularization significantly increases the amount of glucose delivered to islet β-cells, initiating the β-cell maturation in the core and boosting all the glucose-responsive β-cells to be more functional.

**Figure 3.**
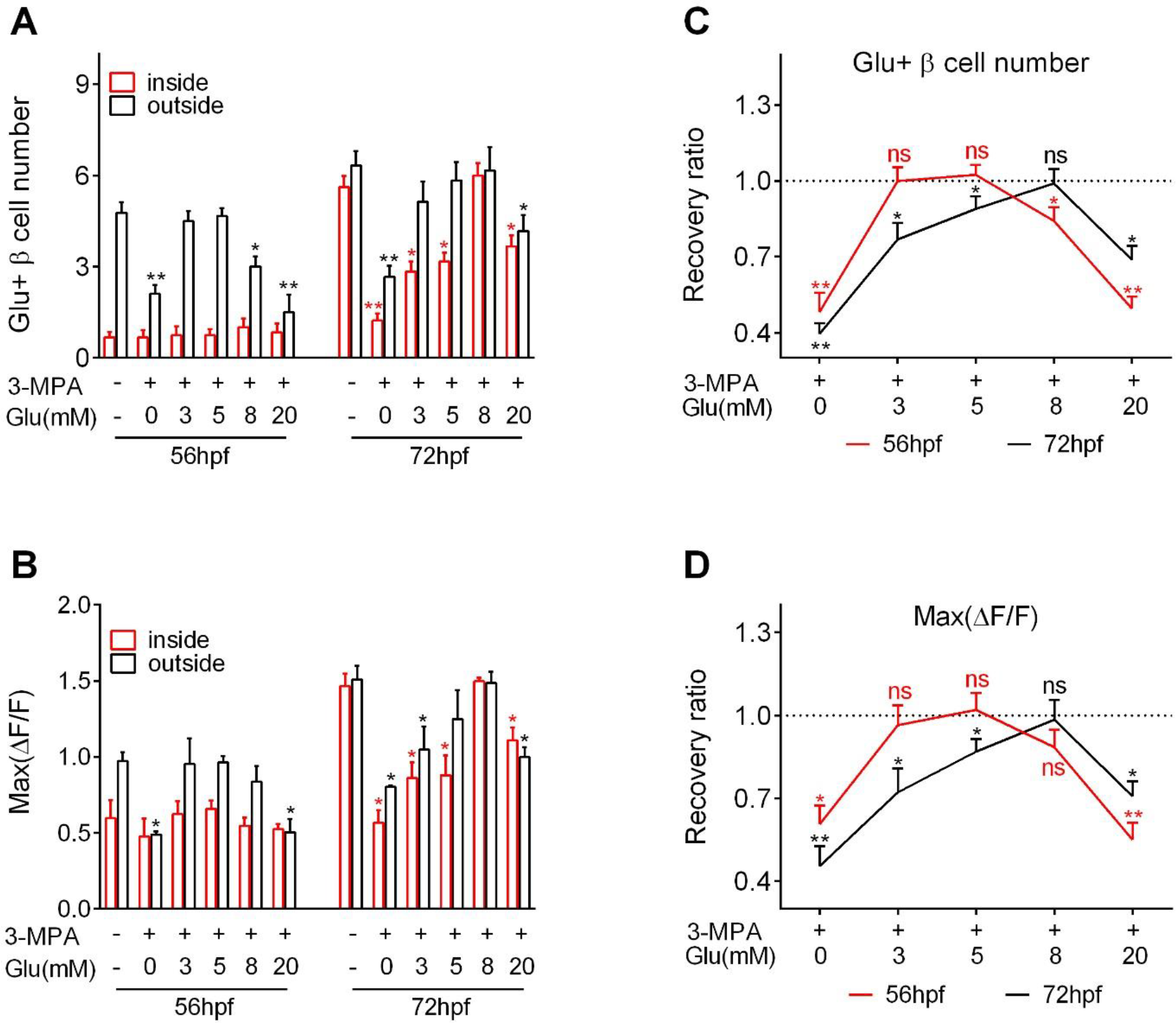
Different concentrations of glucose are required for inducing optimal β-cell maturation at different maturation stages.(*A-B*) Numbers of glucose-responsive β-cells (A) and their maximal Ca^2+^ responses to glucose (*B*) in the mantle and in the core of the islets from embryos treated with 3-MPA and different concentrations of glucose. n = 4-8 embryos for each condition in A and B. \**P*<0.05, \*\**P*<0.01. (*C-D*) The recovery ratios of glucose-responsive β-cells (*C*) and their maximal Ca^2+^ responses to glucose (*D*) by different concentrations of exogenous glucose while the endogenous glucose production was inhibited by 3-MPA. The ratios are represented as the normalized numbers of glucose-responsive β-cells or the normalized maximal amplitudes of the calcium transients relative to those of the control embryos. n = 4-8 embryos for each condition in *C* and D. \**P*<0.05, \*\**P*<0.01; ns, not significant. See also Figure 3-Figure Supplement 1 and Supplemental Movie 5.

### Calcineurin/NFAT activated by glucose drives β cells maturation

Next, we tried to identify the master signalling pathway triggered by glucose. Recently, activation of calcineurin/NFAT by a glucokinase activator was proposed to mediate β-cells development and function in mice (Goodyer et al, 2012a; Heit et al, 2006). To test whether calcineurin/NFAT act as the primary maturation signalling pathway downstream of glucose, we examined the β-cell maturity after incubating the embryos with calcineurin/NFAT inhibitors and activators. From 36 to 48 hpf, either inhibition of calcineurin with FK506 (Goodyer et al, 2012a; Heit, 2007) or suppression of gluconeogenesis with 3-MPA prevented the appearance of the first glucose-responsive β-cell without affecting β-cell proliferation (Figure 4A). In contrast, inhibition of the mTORC pathway with rapamycin abolished β-cell proliferation without affecting its maturation (Figure 4A and B and Supplemental Movie 6). These results suggest that glucose activated calcineurin/NFAT is essential for β-cell maturation initiation.

**Figure 4.**
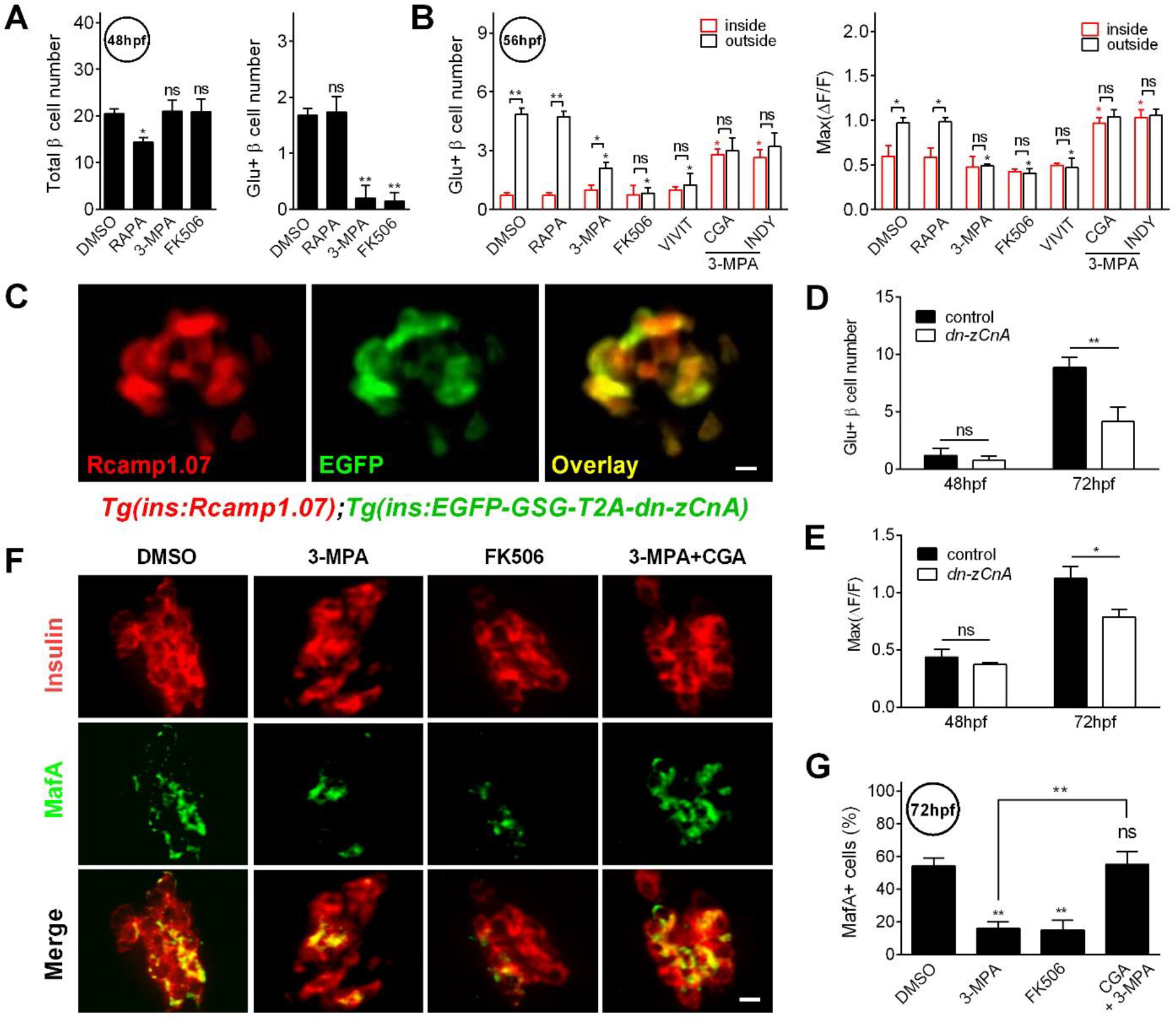
Calcineurin/NFAT acts as the master regulator downstream of glucose to initiate and sustain β-cell maturation. (*A*) Numbers of total β-cells (left) and glucose-responsive β-cells (right) in 48 hpf embryos that had been treated with the indicated reagents. n = 4-8 embryos for each condition. \**P*<0.05, \*\**P*<0.01; ns, not significant. (*B*) Numbers of glucose-responsive β-cells (left) and their maximal amplitudes of Ca^2+^ transients in glucose-responsive β-cells (right) in the mantle and core of the islet in 56 hpf embryos that had been treated with the indicated reagents. n = 5-9 embryos for each condition. \**P*<0.05, \*\**P*<0.01; ns, not significant. (*C*) Representative TPM images of Rcamp1.07 (left), EGFP (middle) and the merged images (right) of the same islet cells in 72 hpf living *Tg(ins:Rcamp1.07);Tg(ins:EGFP-GSG-T2A-dn-zCnA*) embryos. (*D-E*) Numbers of glucose-responsive β-cells (*D*) and their maximal Ca^2+^ responses to glucose (*E*) in the age-matched control and the embryos expressing *dn-zCnA* at 48 and 72 hpf. n = 3-4 embryos for each stage in *D* and *E*. \**P*<0.05, \*\**P*<0.01; ns, not significant. (*F*) Immunofluorescent labelling of insulin and MafA in 72 hpf embryos that had been treated with the indicated reagents. (G) Quantifications of MafA-positive β-cells in 72 hpf embryos that had been treated with the indicated reagents. n = 4-6 embryos for each condition. \*\**P*<0.01; ns, not significant. (Scale bars: 10 μm; scale bars apply to *C* and *F*.) See also Figure 4-Figure Supplement 1 and Supplemental Movie 6.

Inhibiting calcineurin or NFAT (with VIVIT (Demozay et al, 2011)) during the hatching period also significantly reduced the number of glucose-responsive β-cells and their maximal Ca^2+^ response amplitude (Figure 4B and Figure 4-Figure Supplement 1). Interestingly, the β-cells that had been already responsive to glucose before the inhibitor treatment were reverted to less matured states (Figure 2-Figure Supplement 2), suggesting that calcineurin/NFAT continuous activation is required for the maintenance of the maturation state. On the other hand, direct activation of calcineurin with chlorogenic acid (CGA) (Tong et al, 2007) or NFAT with ProINDY (Ogawa et al, 2010) readily rescued the defective functional maturation of β-cells caused by 3-MPA (Figure 4B, Figure 4-Figure Supplement 1 and Supplemental Movie 6). These results suggest a necessary and sufficient role of calcineurin/NFAT in mediating the effect of glucose in initiating, boosting and sustaining β-cell maturation.

To provide further evidence for such a hypothesis, we created and applied a dominant-negative *zebrafish calcineurin A (dn-zCnA)*. As *CnA* is highly conserved with 81% identity in amino acids between zebrafish and human, we constructed *dn-zCnA* based on the strategy of generating dominant-negative *human calcineurin A (dn-hCnA*) (Faure et al, 2007). In *Tg(ins:Rcamp1.07*) genetic background, we generated transient transgenic zebrafish embryos *Tg(ins:EGFP-GSG-T2A-dn-zCnA*), in which the *dn-zCnA* is co-expressed with EGFP with a *GSG-T2A* linker under the drive of the insulin promoter. Under TPM, we observed 75.8% ± 2.5 Rcamp1.07 positive cells co-express EGFP, indicating 70%-80% β-cells express *dn-zCnA* in *Tg(ins:Rcamp1.07*); *Tg(ins:EGFP-GSG-T2A-dnCnA*) double transgenic fish (Figure 4C). When stimulating with 20 mM glucose, we observed a decreased number of glucose-responsive β-cells and smaller maximum Ca^2+^ transients in the responding β-cells in the 72 hpf embryos expressing *dn-zCnA* compared with the age-matched controls (Figure 4D and E). The first glucose-responsive β-cells still appeared at the 48 hpf embryos expressing *dn-zCnA*, but all these glucose-responsive β-cells were EGFP negative, indicative of not expressing *dn-zCnA* in these β-cells. Taken together, these results are consistent with the results of pharmacological treatments with calcineurin/NFAT inhibitors, providing convincing evidence that calcineurin/NFAT drives β-cell maturation.

Finally, inhibiting calcineurin or suppressing gluconeogenesis severely reduced the number of MafA-positive β-cells at 72 hpf to a similar extent. Direct activating calcineurin, on the other hand, fully rescued the number of MafA-positive β-cells in 3-MPA treated fish (Figure 4F and G). Thus, glucose activates the calcineurin/NFAT signalling pathway to initiate, boost, and sustain the whole process of β-cell maturation, possibly via upregulation of MafA.

### Direct activating calcineurin promotes neonatal β-cells towards optimal maturity in isolated mouse islets in vitro

The failure for the core β-cells to mature in the *cloche* mutants and 2,3-BDM-treated embryos could be due to reduced glucose delivery to the islet centre. Indeed, in isolated mouse islets that had no microcirculation, we found a gradually reduced glucose transported into β-cells of the inner layers (Figure 5A-C). To explore whether the requirement of optimal glucose for β-cell maturation could be bypassed *in vitro,* we tried to activate calcineurin/NFAT to promote β-cell maturation in isolated neonatal mouse islets. As we have known, the β-cells of neonatal mouse islets respond poorly to glucose and need more than one week to mature *in vivo* (Blum et al, 2012b). We cultured the neonatal islets from postnatal day 0 (P0) mice for 3 days in media supplemented with different concentrations of glucose, in the absence or presence of the calcineurin activator CGA, and evaluated their maturity according to the glucose stimulation index (GSI). Among the different glucose concentrations tested, 11 mM glucose in the culture media was the best in transforming neonatal β-cells in islets to be more glucose-responsive (GSI = 13.0 ± 1.3, Figure 5D and E). Supporting a conserved role of calcineurin in mouse β-cell maturation, combining CGA in the culture media greatly enhanced the GSIs of islets cultured at a glucose concentration ranging from 5.6 mM to 11 mM. Neonatal islets cultured at 11 mM glucose and 56.48 μM CGA exhibited a GSI of 26.5 ± 4.8 (n = 12), similar to that of adult mouse islets cultured at 11 mM glucose (27.3 ±3.4, n = 4, p = 0.79) (Figure 5D and E). On the other hand, calcineurin inhibitor FK506 notably reduced the GSI of the neonatal islets cultured at 11 mM glucose (Figure 5D and E). As compared to more than one week needed for P0 β-cells to mature *in vivo* (Blum et al, 2012b), our results indicate a greatly accelerated maturation process *in vitro* by using small molecule activators of calcineurin/NFAT, which may also be used to push stem-cell derived immature β-cells towards optimal maturity *in vitro*.

**Figure 5.**
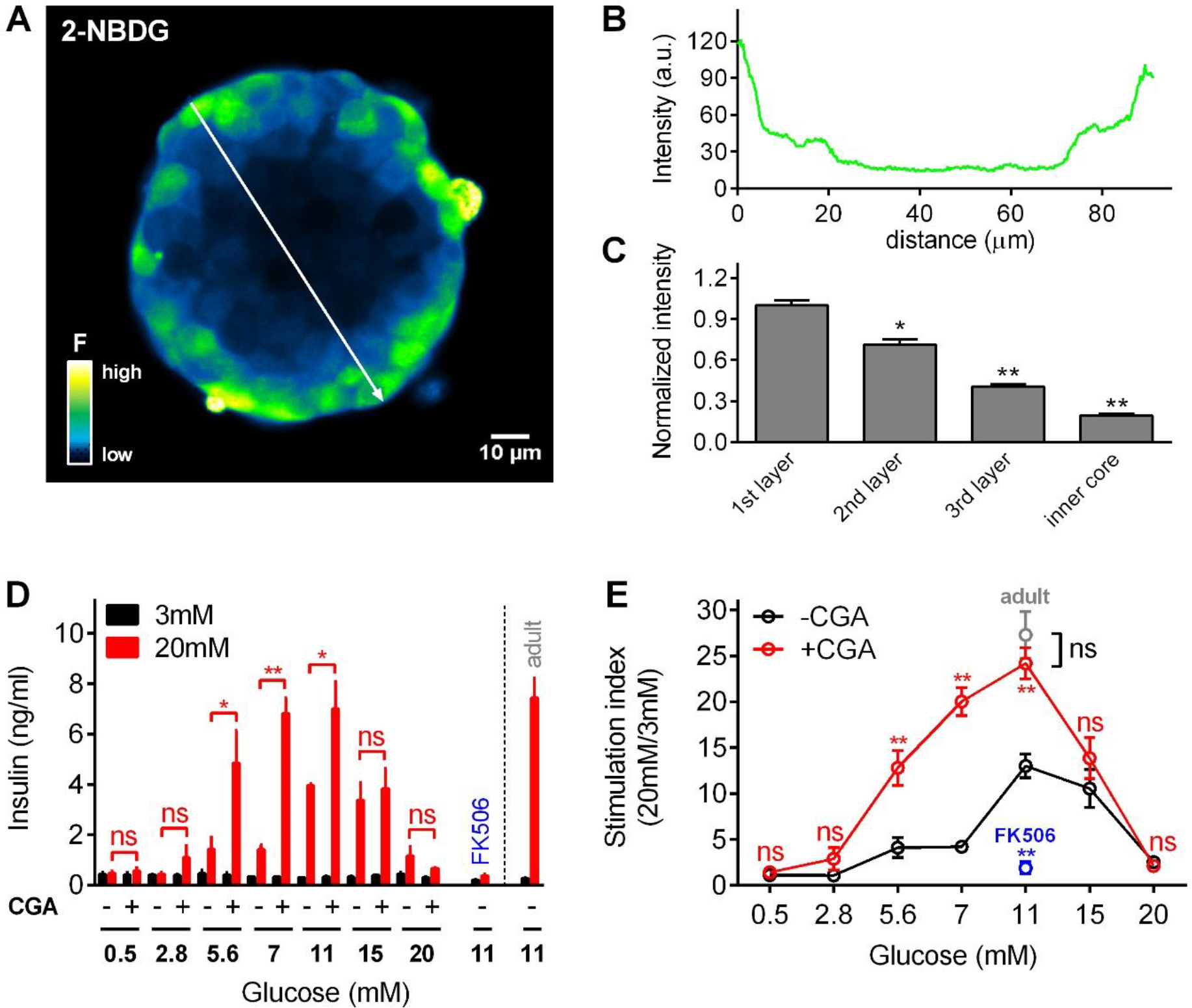
Direct activating calcineurin readily pushes neonatal mouse β-cells in isolated islets towards optimal maturation *in vitro*. (*A*) Significantly reduced glucose uptake in the core of the islets *in vitro* represented as the TPM image of an islet pre-incubated with 7mM 2-NBDG (a fluorescent D-glucose analog) for 10 min. (*B*) Profile plot of fluorescent intensity versus distance indicated in A (white line with arrow). (*C*) Normalized fluorescent intensity of β-cells in each layer of the islet in A. \**P*<0.05, \*\**P*<0.01. (*D*) Glucose-stimulated insulin secretion from neonatal mouse islets stimulated with low and high glucose, after isolation from P0 mice and culturing *in vitro* for 3 days in different media. Insulin secretion of adult islets from 8-week-old mice after 3 days of culturing in media with 11 mM glucose is also shown. n = 4-10 experiments for each condition. \**P*<0.05, \*\**P*<0.01; ns, not significant. (*E*) GSIs of the neonatal islets and the adult islets described in D. n = 4-10 experiments for each condition. \*\**P*<0.01; ns, not significant.

## Discussion

Here, by taking advantage of the advanced microscopy 2P3A-DSLM technique, we were able to resolve individual β-cells and monitor their functions in an intact living organism. Spatiotemporally, we characterized the maturation of β-cells *in vivo* as two waves that are differentially affected by islet microcirculation: before islet vascularization, locally synthesized glucose initiates the early β-cell maturation from the islet mantle only; islet vascularization significantly increases the amount of glucose delivered to islet β-cells, initiating maturation of β-cells in the islet core and pushing all glucose-responsive β-cells towards further maturity (Figure 6).

**Figure 6.**
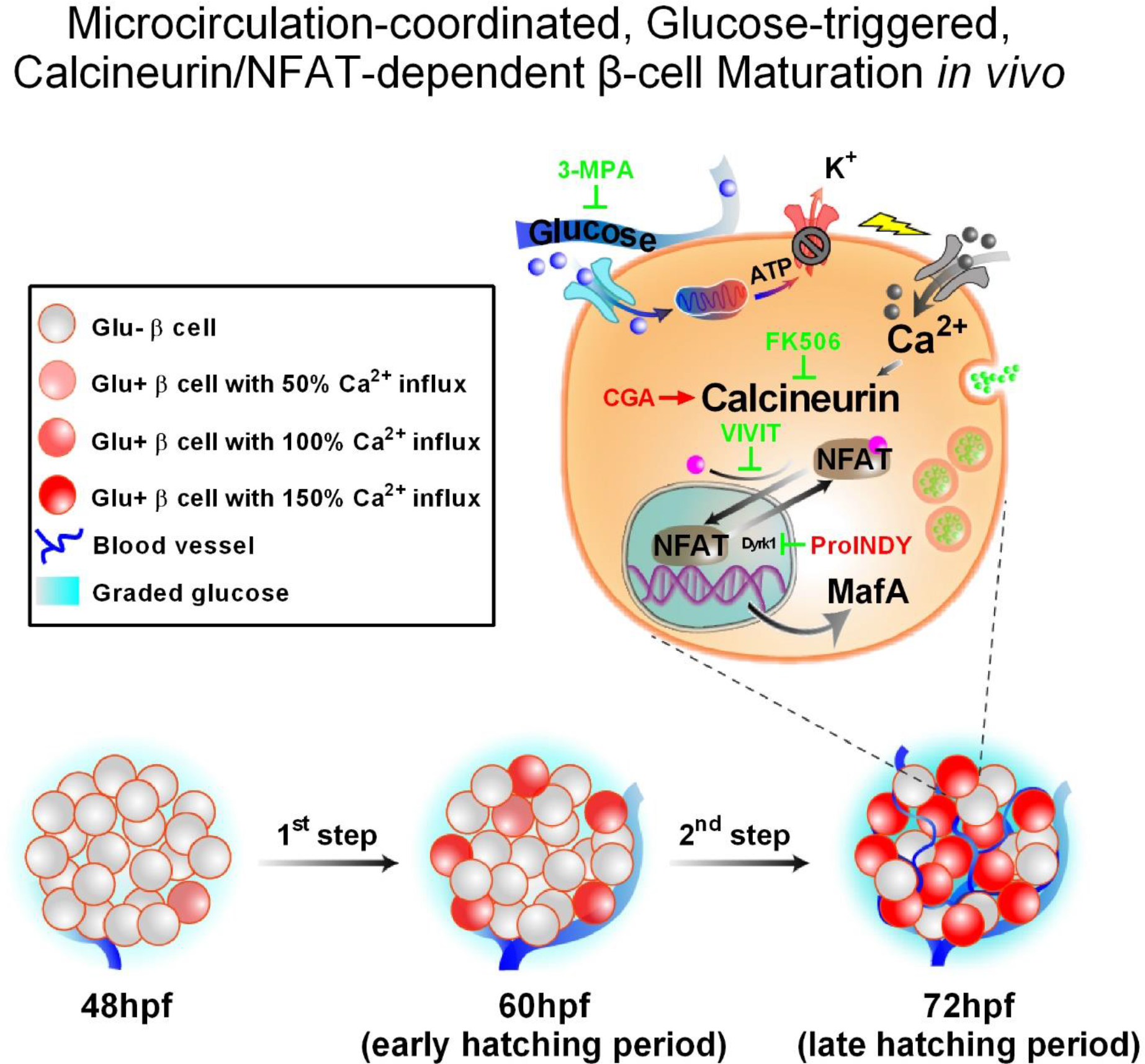
An inter-species conserved model of microcirculation-coordinated, glucose-triggered, calcineurin/NFAT-dependent β-cell maturation. *in vivo*. At approximately 48 hpf, locally synthesized glucose (∼3 mM) initiates glucose responsiveness of β-cells exclusively in the islet mantle. After 60 hpf, islet vascularization enables the efficient delivery of glucose through the blood circulation to the islet core to trigger the internal cells to mature. Both the mantle and the core β-cells sense a more than 2-fold increase in glucose (∼8 mM), which pushes them to further maturity. Maturation of all β-cells in the islet depends on the glucose-activated calcineurin/NFAT pathway, the master regulators of MafA expression and β-cell function acquisition.

With classical methods such as immunofluorescence microscopy to observe the expression of maturation markers in fixed samples or ELISA to measure the GSI, β-cells maturation is usually evaluated as a whole. Whether all β-cells in the islets mature at a synchronized pace remains mysterious. By longitudinal imaging β-cell function *in vivo,* we have discovered the apparent heterogeneity of the maturation process inside the islets. In addition, our data also suggests that the dosage effect of glucose on maturation initiation is different from that on function boosting. Suppression of the *in vivo* glucose production with 3-MPA at the early hatching period did not prevent all the mantle β-cells from entering maturation initiation, but limited their glucose-evoked Ca^2+^ responses to the lowest level as those in the core β-cells (Figure 4B). Therefore, the glucose concentrations needed for the conversion of glucose-nonresponsive β-cells to glucose-responsive β-cells may be lower than those required for further boosting β-cell function thereafter. Consistently, the glucose concentration needed for the further maturation of β-cells from 55 hpf to 72 hpf increases significantly (Figure 3), reinforcing the concept of a delicately regulated glucose concentration for different steps of β-cell maturation. As zebrafish and mammals share many similarities during pancreatic islet development (Supplemental Table 1), an increasing dosage requirement of glucose for β-cells maturation *in vivo* is very likely to be a conserved mechanism. Consistent with this hypothesis, the plasma glucose level in fetal rodents is relatively low, but starts to increase just before birth and progressively reaches a plateau (above 4 mM) after P2 (Rozzo et al, 2009). From P6 to P20, the plasma glucose further gradually increases from 6 mM to 12 mM (Aguayo-Mazzucato et al, 2006). This period is just accompanied with islet vascularization establishment and glucose-responsive β-cells appear and further maturation (Hole et al, 1988; Reinert et al, 2014; Rozzo et al, 2009). Therefore, β-cell maturation in rodents may also require a dynamically increased glucose concentration, coordinated by islet vascularization.

Multiple signalling pathways are known to be activated by glucose to regulate gene transcriptions. These “master switches” include calcineurin/NFAT (Lawrence et al, 2002), the AMP-activated protein kinase, carbohydrate response element-binding protein (Iizuka & Horikawa, 2008; Vaulont et al, 2000), and the hexosamine biosynthetic pathway (Vanderford et al, 2007). Although implication of calcineurin/NFAT in β-cell maturation has been proposed in mice (Goodyer et al, 2012a; Heit et al, 2006), their critical role on the β-cell proliferation mask their effect on the β-cell maturation. For instance, β-cell specific *calcineurin b1* knock-out mice (*RIP-Cre; Cnb1^Δ/f^*) did not exhibit any phenotypes until 8-week after birth when β-cell mass starts to reduce (Heit et al, 2006). In contrast, pancreatic endocrine progenitor cell specific *calcineurin b1* knock-out mice (*Ngn3-Cre; Cnb1^Δ/lax^*) developed glucose intolerance and diabetes at P20, accompanied with a more than 80% reduction in β-cell mass at P26 (Goodyer et al, 2012a). These data strongly support a critical role of calcineurin/NFAT in β-cell precursor’s differentiation and β-cell proliferation, but obscure its role in β-cell maturation. The failure to establish a solid link between calcineurin/NFAT and β-cell maturation may explain why manipulation of this pathway has never been tried to enhance stem cell-derived β-like cells maturation *in vitro*. By imaging β-cell function acquisition and enhancement in zebrafish *in vivo*, we demonstrate that calcineurin/NFAT, downstream of increasing glucose concentrations delivered by the blood circulation, is essential for the initiation, boosting and maintenance of β-cell maturation. The mechanism is inter-species conserved, as activating calcineurin/NFAT could also bypass the deficit of microcirculation in isolated mouse islets, accelerate the *ex vivo* maturation of neonatal mouse β-cells (Figure 5D and E). Currently, different research groups use a variety of glucose concentrations, ranged from 2.8 mM to 20 mM, in the last stage to push β-like cells derived from stem cells *in vitro* to be more functional competent, as nobody knows the ideal condition for the β-cell maturation (Bruin et al, 2015). Obviously, blood vessels are absent from these β-like cell clusters, either. Based on our data, we argue that the lack of spatiotemporal precise glucose delivery to all β-like cells may contribute significantly to their limited glucose responsiveness. On this regard, direct activation of calcineurin/NFAT with small molecules may confer a previously unexplored strategy to push these β-like cell clusters toward maximal maturity *in vitro*.

Future imaging of panoramic β-cell mass and function *in vivo* in zebrafish with our system may unravel mechanisms of other important but obscure processes in islet biology, such as regeneration, cell identity or functional changes during disease progression. Understanding these principals may shed light on the ultimate cell replacement therapy to treat diabetes.

## Materials and Methods

### Transgenic zebrafish generation

The Tg(*ins:Rcamp1.07*) reporter zebrafish line was generated using meganuclease-mediated transgenesis as previously described (Soroldoni et al, 2009). Briefly, zebrafish BAC_CH211_69I14 (BACPAC Resources Center), which contains the zebrafish *insulin* gene, was modified using Red/ET recombineering technology to replace the coding sequence of insulin with *Rcamp1.07* (Fu et al, 2010). Rcamp1.07-tagged BAC underwent a second round of recombineering for the subcloning of the modified chromosomal locus into a plasmid backbone containing two I-SceI meganuclease sites. The following primers were used for BAC modification:

Forward primer: 5’ ATGTTTTTGATTGACAGAGATTGTATGTGTGTGTTTGTGTCAGTGTGACCCGCCACCATGGGTTCTCATC.
Reverse primer: 5’ CCTGTGTGCAAACAGGTGTTTCTGGCATCGGCGGTGGTCAAATCTCTTCAGGCAGATCGTCAGTCAG.
And for subcloning:
Forward primer: 5’ AGTCCATTAAATAATATCTTGTAGAATTATGTTTTTAAAAAGTACCAATGCCGTAGGGATAACAGGGTAATTTAAGC.
Reverse primer: 5’ CAACTTTTTCACAAACACTGACCAAAACAAGCTACATGTTTTAGAGGCATTAGGGATAACAGGGTAATTGCACTG.

The resulting constructs were co-injected with I-SceI meganuclease (Roche) at a DNA concentration of 100 ng/μl into one-cell stage zebrafish embryos. These F0 embryos were screened for the transient expression of Rcamp1.07 in the pancreatic islet at 48 hpf using a fluorescence stereomicroscope. The positive F0 founders were raised to adults and were screened by visual inspection of their F1 progenies from outcrossing with the wild-type AB strain. Based on the intensity of the fluorescence signal, one founder was selected, and subsequent generations were propagated and expanded.

The *Tg(ins:EGFP-GSG-T2A-dn-zCnA*) zebrafish line was generated using Tol2 transposase RNA-mediated transgenesis as previously described. Briefly, *dn-zCnA* was constructed by deleting the autoinhibitory and the calmodulin-binding domains through introducing a stop codon at the N396 amino acid and by mutating the histidine at the position 152, a phosphatase-active site, to glutamine. The GSG-T2A peptide was used to separating EGFP and dn-zCnA elements. The EGFP-GSG-T2A-dn-zCnA fragment was cloned downstream of 2.1 kb of the proximal insulin promoter and into a Tol2 plasmid by ClonExpress system. The final construct was injected along with Tol2 transposase RNA into Tg(*ins:Rcamp1.07*) eggs to generate mosaic Tg(*ins:EGFP-GSG-T2A-dn-zCnA*); Tg(*ins:Rcamp1.07*) F0 fish for imaging analysis.

### Zebrafish care and handling

The wild-type AB strain and transgenic fish were maintained and handled according to the institutional guidelines of animal usage and maintenance of Peking University. *Tg(ins:EGFP*) fish were from Dr. Lin Shuo at UCLA; *Tg(flk1:mCherry*) fish were from Dr. Zhang Bo at PKU; *Tg(flk1:GFP*) fish were from Dr. Chen Jau-Nian at UCLA. *Tg(ins:Rcamp1.07*) was crossed with heterozygous *cloche^m39^* to obtain *Tg(ins:Rcamp1.07);cloche ^m39^/+* zebrafish. *Tg(ins:Rcamp1.07);cloche ^m39^/ cloche ^m39^* embryos from the crossing of *Tg(ins:Rcamp1.07);cloche ^m39^/+* with *cloche ^m39^/+* fish were used for the imaging experiments. For imaging, 0.002% phenylthiourea (PTU, Sigma) was added at 12 hpf to prevent pigment synthesis. Prior to live imaging, embryos were anaesthetized with 0.01% tricaine (Sigma).

Pharmacological treatment. Heterozygous *Tg(ins:Rcamp1.07*) embryos were used for pharmacological treatment and imaging analysis. All chemicals were prepared as high-concentration stocks and were diluted in E3 medium to final concentrations for treatment, which were carefully selected to be non-toxic and effective. To test the pharmacological effect of the chemicals on the functional maturation of pancreatic β cells, the embryos were treated with each chemical for 9 hours, either from 44 to 53 hpf or from 60 to 69 hpf; the embryos then recovered for 3 h after E3 medium was used to completely wash out the chemical, and imaging analysis was carried out at 56 hpf or at 72 hpf, respectively. For the pharmacological treatment of zebrafish embryos, 10 mM 2,3-butanedione monoxime (2,3-BDM, Sigma) was used to block blood circulation; 3 mM 3-mercaptopicolinic acid (3-MPA, Santa Cruz) was used to inhibit endogenous glucose production; 141.2 μM chlorogenic acid (CGA, Sigma) or 10 μM FK506 (Invivogen) was used to activate or inhibit calcineurin, respectively; and 2.5 ProINDY (Tocris) or 20 VIVIT peptide (Tocris) was used to activate or inhibit NFAT, respectively.

### Wide field and 2P3A-DSLM imaging

For the wide field time-lapse imaging, the images were captured using an Olympus inverted fluorescence microscope with a 10×/0.4 objective. D-Glucose stock solution was added to the E3 medium for a final concentration of 20 mM during stimulation. The images were collected with MetaMorph software and analysed with Fiji software.

High-resolution images of pancreatic β-cells and blood vessels in live zebrafish embryos were captured with 2P3A-DSLM equipped with two 40×/0.8 water lenses as previously described (Zong et al, 2015). Briefly, the anaesthetized embryos were embedded in a 1% ultra-pure agarose (Invitrogen) cylinder and then immersed in a homemade chamber with E3 medium containing 0.01% tricaine. D-glucose stock was added to the E3 medium to reach a final concentration of 20 mM to stimulate the fish embryos. For the 2D time-lapse imaging experiments that were used for statistical analysis, the islet was optically sectioned into 5-6 layers to ensure that the calcium transients of all β-cells within the islet were recorded. For fast volumetric imaging and reconstruction of calcium transients within the whole islet, the islet was optically sectioned into 25 layers. Each layer was captured 5 times with an 8-ms exposure time and was averaged as one single image. Images were collected by the HCImage software (Hamamatsu) and processed with R-L deconvolution by Fiji software. The volumetric calcium transients were reconstructed with Amira software (FEI).

### Comparison among 1P-SPIM, TPM and 2P3A-DSLM in 3D imaging of islet in vivo

For 1P-SPIM imaging, we used a homemade SPIM setup equipped with a 40×/0.8 water lens. For TPM imaging, we used a fast resonant-scanned TPM equipped with a 40×/0.8 water lens. The same 72 hpf zebrafish samples were sequentially imaged with 1P-SPIM, TPM and 2P3A-DSLM. Under each configuration, the whole islet was optically sectioned by 100 planes (z-step: 500 nm) with an exposure time of 150 ms per frame. Images were collected by HCImage software and processed with R-L deconvolution by Fiji software.

### Whole-mount immunofluorescence and confocal imaging

The embryos were fixed in 4% paraformaldehyde (PFA, AppliChem) at 4 °C overnight. After the PFA was washed away, the embryos were dehydrated using a series of methanol (25%, 50%, 75% and 100%) and stored at -20°C. When necessary, the embryos were rehydrated using a series of methanol (100%, 75%, 50% and 25%), permeabilized in Proteinase K (10 μg/ml, TransGen), washed in PBST (PBS + 0.1% Tween-20) and then fixed again in 4% PFA at room temperature (RT). After being blocked with PBST containing 0.2% bovine serum albumin (BSA, AppliChem) and 5% fetal bovine serum (Gibco) for 1 h at RT, the embryos were incubated with primary antibodies overnight at 4°C. Then, secondary antibodies were applied at 4 °C overnight after thorough washing. The primary antibodies included monoclonal rat anti-mCherry antibody (1:200, Thermo, used to detect Rcamp1.07), polyclonal guinea pig anti-insulin antibody (1:200, Dako) and polyclonal rabbit anti-MafA antibody (1: 50, Sigma). The secondary antibodies were Alexa Fluor 568 goat anti-rat IgG (1: 500, Thermo), DyLight 488 goat anti-guinea pig IgG (1: 500, Thermo), DyLight 550 goat anti-guinea pig IgG (1: 500, Thermo) and DyLight 488 goat anti-rabbit IgG (1: 500, Thermo). Before imaging, the samples were dehydrated in 100% methanol and mounted in mounting solution containing one volume of benzyl alcohol (Sigma) and two volumes of benzyl benzoate (Sigma) with their right side facing the coverslips. The images were collected by MetaMorph software using an Olympus spinning-disc confocal microscope and processed by Fiji software.

### Embedding and cryostat sectioning

The embryos were fixed in 4% PFA at 4 °C overnight. After being washed, the embryos were dehydrated in 30% sucrose (Sigma) and then transferred to embedding chambers filled with OCT compound (Tissue Tek^®^). After embedding, the samples were frozen in liquid nitrogen as soon as possible. Sectioning was performed using a Leica CM1900 Cryostat set to 10 μm thickness and a −25 °C chamber temperature. The sections were collected and kept at -20 °C in a sealed slide box. Immunofluorescence staining was performed as described above. For nuclear staining, the samples were incubated with 2 μg/ml DAPI (Solarbio) for 10 minutes. After an extensive wash, the samples were mounted in 80% glycerol (Sigma) and imaged using an Olympus spinning-disc confocal microscope.

### Uptake of the fluorescent D-glucose analogue by β-cells in mouse islets

For this study, 8-week-old mouse islets were isolated and cultured in RPMI 1640 medium containing 7 mM glucose for 2 days (Wang et al, 2016). Before imaging, the islets were washed twice with Krebs (KRB) buffer (129 mM NaCl, 4.7 mM KCl, 1 mM CaCl_2_ • 2H_2_O, 1.2 mM MgSO_4_, 1.2 mM KH_2_PO_4_, 5 mM NaHCO_3_, 10 mM HEPES, 0.5% BSA) and then pre-incubated in 7 mM 2-(N-(7-nitrobenz-2-oxa-1,3-diazol-4-yl)Amino)-2-deoxyglucose (2-NBDG) dissolved in KRB buffer for 10 min at 37°C. After thoroughly washing, z-stack images of the islets were captured with a TPM (Zeiss 710). The 2-NBDG signal was excited at 920 nm and collected between 510 nm and 540 nm.

### In vitro culture of mouse islets and measurement of glucose-stimulated insulin secretion

Adult islets from 8-week-old mice were isolated as previously described (Wang et al, 2016). For P0 mouse islet isolation, pancreata were dissected directly without perfusion and digested with 0.5 mg/ml Collagenase P (Roche). The isolated islets were cultured for 3 days in RPMI1640 media containing different concentrations of glucose (0.5 mM, 2.8 mM, 5.6 mM, 7 mM, 11 mM, 15 mM and 20 mM) combined or not combined with CGA (56.48 μM). The culture medium was changed every day. 10 islets with similar sizes were selected and pre-incubated in KRB buffer for 3 hours at 37°C in a 5% CO_2_ incubator. The islets were then transferred into low glucose (3 mM) KRB buffer and incubated for 1 hour at 37°C, 5% CO_2_. The supernatant was collected for measuring basal insulin secretion. The same islets were further transferred into high glucose (20 mM) KRB buffer for 1 hour incubation at 37°C, 5% CO_2_. The supernatant was collected, stored at -20°C and later the insulin content was measured with the Rat/Mouse insulin ELISA kit (Millipore).

### Statistical analysis

All data were analysed using GraphPad Prism 6 software. Average results were displayed as the mean values ± SEM. Statistical significance was evaluated using either Student’s t-test for single Gaussian-distributed datasets or Mann-Whitney rank sum test for non-single Gaussian-distributed datasets. The asterisks ^*^ and ^**^ denote statistical significance with *p* values less than 0.05 or 0.01, respectively.

### Study approval

Generation of the transgenic zebrafish lines, *in vivo* imaging of the living zebrafish embryos, and all the other experiments about zebrafish and mouse islets were approved by the IACUC of Peking University.

## Acknowledgments

We thank Dr. Lin Shuo, Dr. Zhang Bo, Dr. Chen Jau-Nian and Dr. Liu Feng for sharing the fish lines with us. We thank Dr. Liao Bo-Kai and Ms. Meng Liying for technique consultant and assistant. This work was supported by grants from the National Science Foundation of China (81222020, 31221002, 31327901, 31570839, 3142800018 and 31301186), the Natural Science Foundation of Beijing Municipality (7121008, 7152079), the Major State Basic Research Program of P.R. China (2013CB531200), and the National Key Technology R&D Program (SQ2011SF11B01041).

## Author contributions

Y. L. conceived the project. L.C. and Y.L. directed the study. J.Z., L.C. and Y.L. designed the research. J.Z., W.Z., Y.W., J.S., D.G., Y.Z., R.W., F.N., X.W., X.Z., A.W., Y.Z., and Y.L. performed the experiments. J.X. provided critical fish lines and fish facility. J.Z., L.C. and Y.L. analyzed the data and wrote the manuscript.

## Figure Legends

**Figure 1-Figure Supplement 1.**
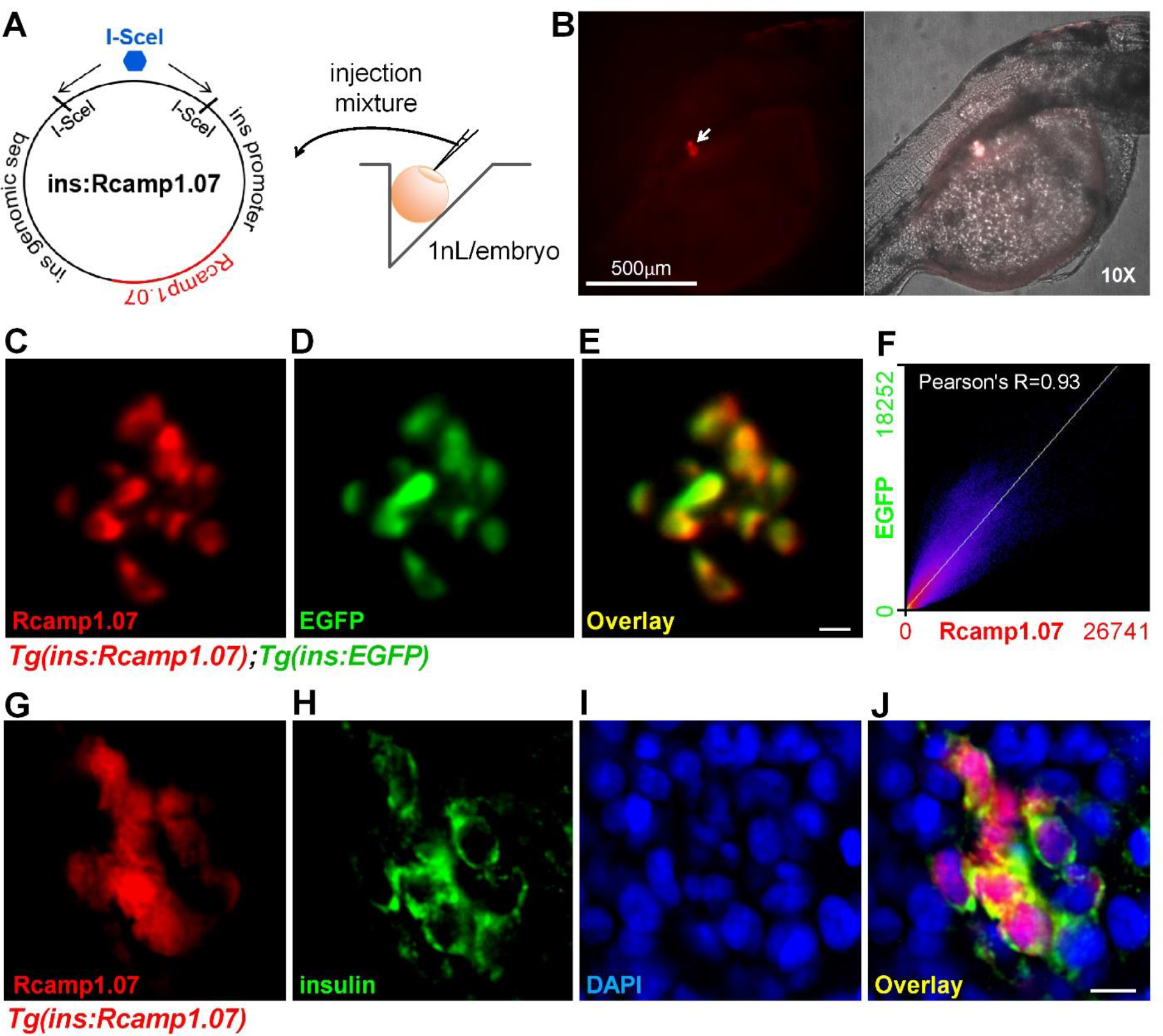
Specific expression of Rcamp1.07 in pancreatic β-cells in *Tg(ins:Rcamp1.07*) zebrafish. (*A*) An illustration shows the protocol for generating the *Tg(ins:Rcamp1.07)* fish line. (*B) Tg(ins:Rcamp1.07*)-positive fish with Rcamp1.07 fluorescent signals in the islet region under a wide field microscope. (*C-E*) Representative dual-colour 2P3A-DSLM images of Rcamp1.07 (*C*), EGFP (*D*) and the merged images (*E*) of the same islet cells in 72 hpf living *Tg(ins:Rcamp1.07);Tg(ins:EGFP)* embryos. (*F*) The co-localization of Rcamp1.07 and EGFP was analysed by Mander’s intensity correlation (Fiji plugin), and the Pearson's R value was 0.93. (*G-J*) Representative confocal images of immunofluorescently labelled Rcamp1.07 (G), insulin (H), DAPI labelling (*I*) and the merged images (*J*) of the same islet cells in 72 hpf *Tg(ins:Rcamp1.07*) embryos. (Scale bars: 10 μm; scale bars apply to *C-E* and *G-J*.).

**Figure 1-Figure Supplement 2.**
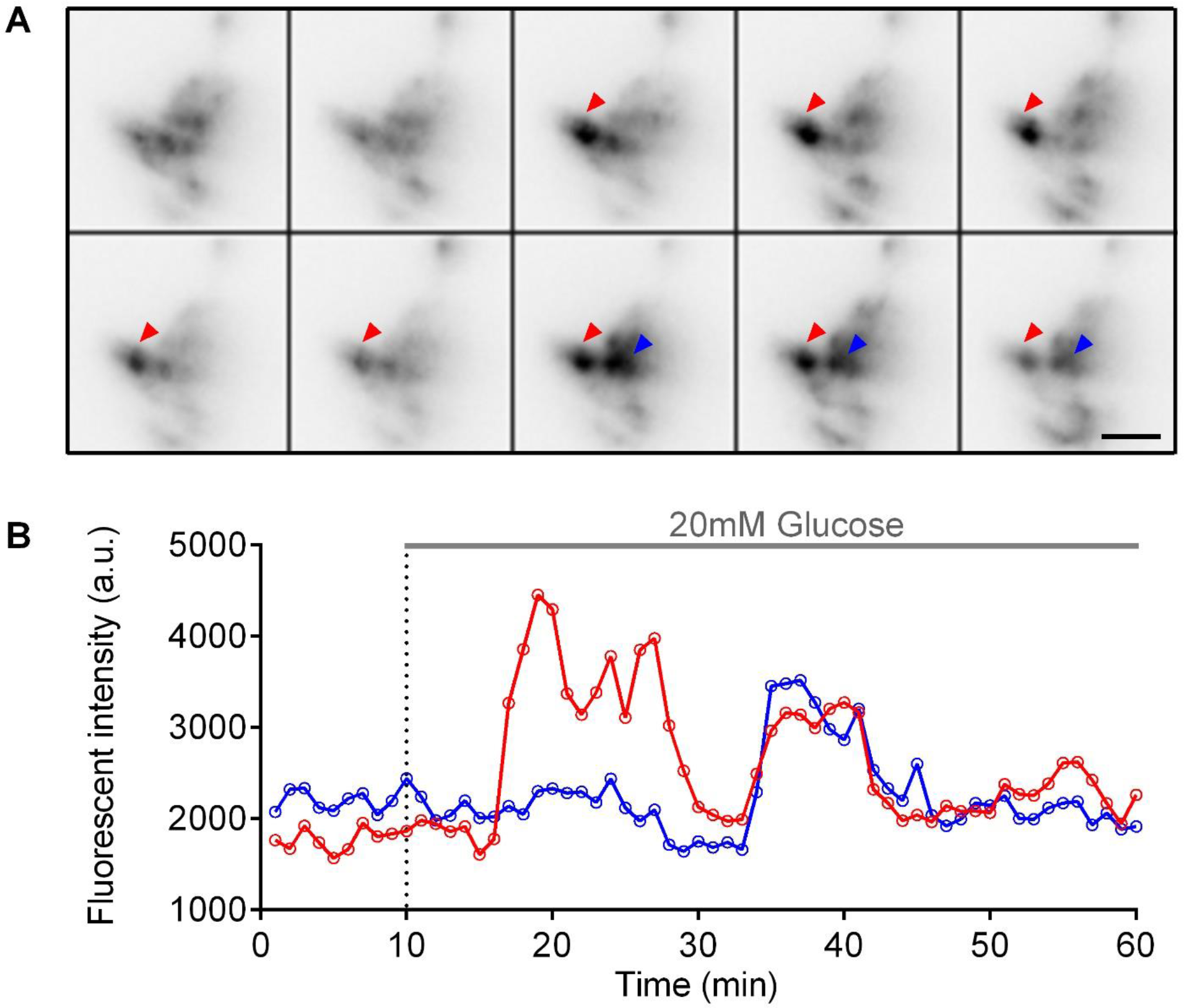
Visualization of glucose-stimulated calcium transients in β-cells in living *Tg(ins:Rcamp1.07*) embryos under a wide field microscope. *Tg(ins:Rcamp1.07*) embryos were embedded in a 1% ultra-pure agarose cylinder and immersed in a 35 mm plastic chamber filled with E3 medium containing 0.01% tricaine. Images were captured once every minute with a 150 ms exposure time. At the 10 min time point, 20 mM glucose was added. (*A*) Representative images of Rcamp1.07 in the pancreatic islet of a 72 hpf live *Tg(ins:Rcamp1.07*) embryo after stimulation with 20 mM glucose. Montages started from 3 min after glucose application and are shown at 3-min intervals. Arrows indicate two representative regions within the islet. (*B*) Time courses of calcium transients from the regions marked in A. (Scale bar: 50 μm.)

**Figure 1-Figure Supplement 3.**
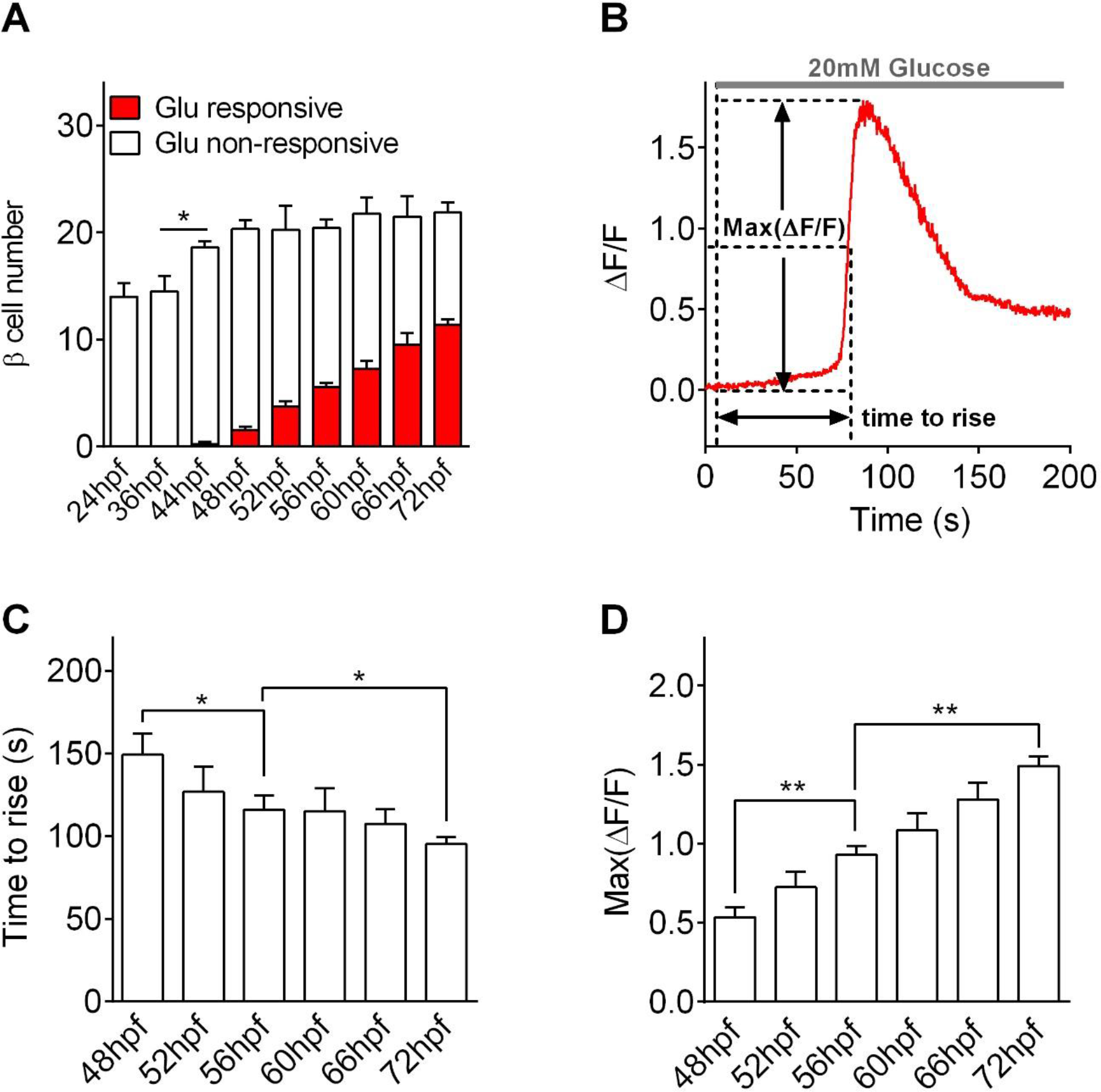
Quantification of glucose-responsive β-cells and evaluation of their maturation states. (*A*) Quantification of glucose-responsive β-cells at different stages from 24 to 72 hpf. \**P*<0.05. (*B*) An illustration of the two parameters describing the kinetics of glucose-stimulated Ca^2+^ transients: the maximum amplitude (Max ΔF/F) and the speed of the Ca^2+^ response (time to rise); the latter was defined as the delay between the time of glucose application and the time when the increase in cytoplasmic Ca^2+^ reached half of the Max ΔF/F. (*C*) Average speeds (time to rise) of the Ca^2+^ responses in glucose-responsive β cells from 48 to 72 hpf. \**P*<0.05. (*D*) Average maximum amplitudes of Ca^2+^ transients in glucose-responsive β-cells from 48 to 72 hpf. \*\**P*<0.01.

**Figure 2-Figure Supplement 1.**
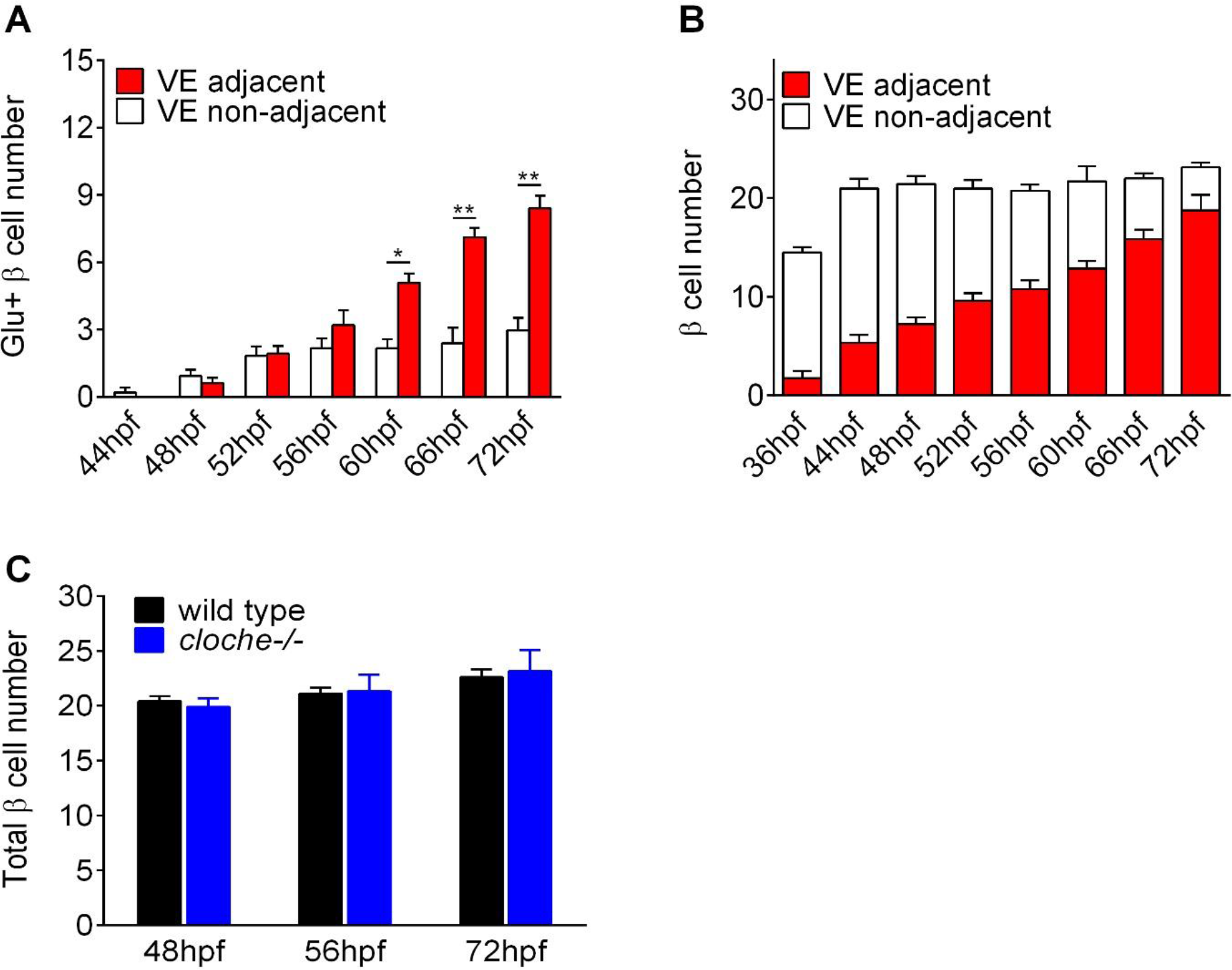
Islet vascularization does not affect the total β-cell number, but correlates with the acquisition of glucose-responsiveness of β-cells. (*A*)Quantification of glucose-responsive β-cells adjacent or non-adjacent to blood vessels from 44 to 72 hpf. n = 4-7 embryos for each condition. (*B*) Quantification of pancreatic β-cells adjacent or non-adjacent to VE cells from 36 to 72 hpf. n = 8 embryos for each condition. \**P*<0.05, \*\**P*<0.01. (*C*) Quantification of total β-cells in wild-type or *cloche*^−/−^ embryos at 48, 56, and 72 hpf. n = 3-4 embryos for each condition.

**Figure 3-Figure Supplement 1.**
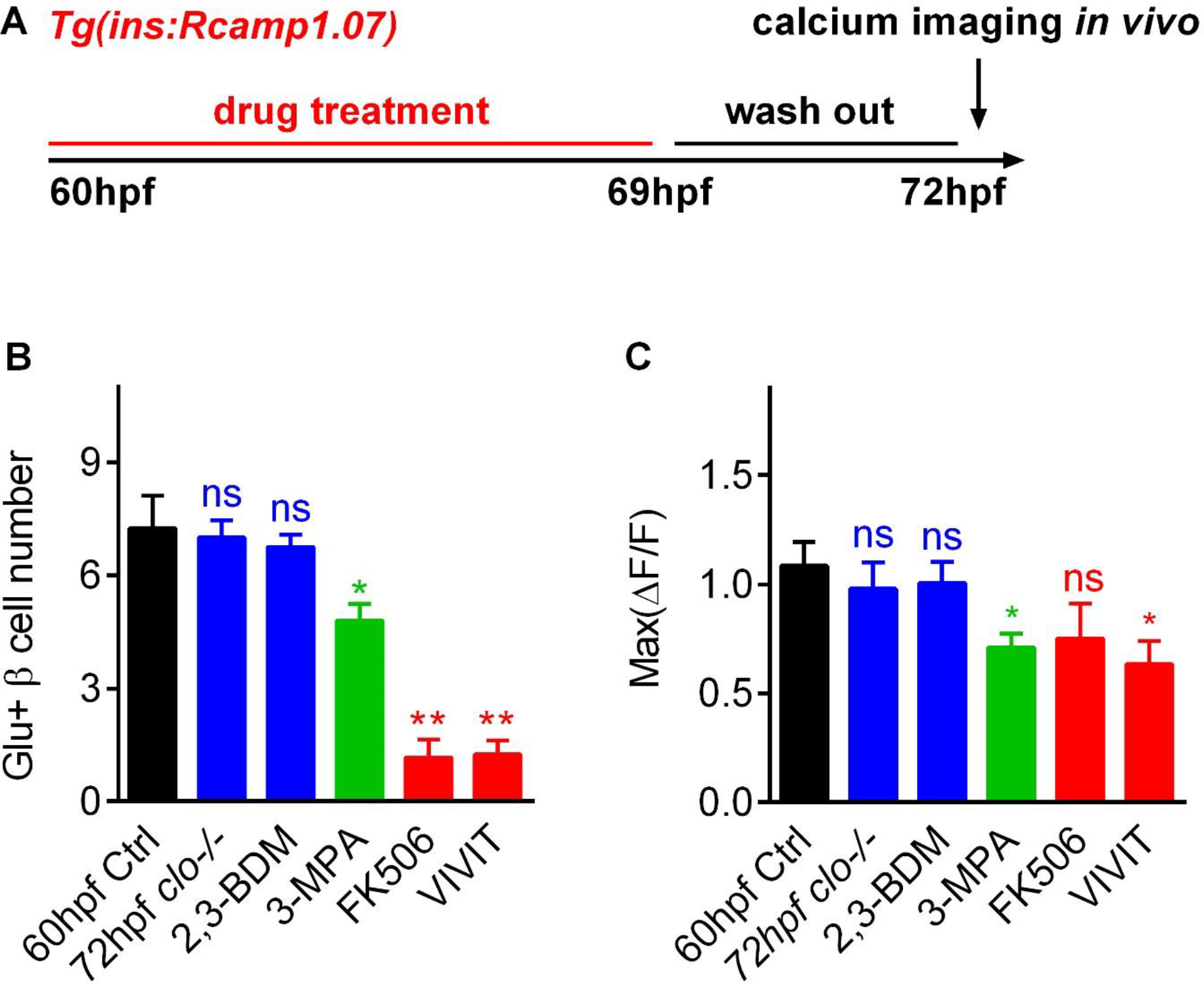
Blocking blood circulation at the late hatching period stops β-cells at a functional status equivalent to 60 hpf embryos, whereas inhibiting the glucose/calcineurin/NFAT pathway severely reverts glucose-responsive β-cells to less matured states. (*A*) The experimental design for *B* and *C* in which *Tg(ins:Rcamp1.07*) embryos at 60 hpf were treated with different reagents for 9 h and then washed in the normal E3 medium for 3 h before imaging. (*B-C*) Quantification of glucose-responsive β-cells (*B*) and their maximal ΔF/F (*C*) in 60 hpf controls (n = 4), 72 hpf *cloche^-/-^* mutants (n = 4) and 72 hpf embryos that had been treated with 2,3-BDM (n = 4), 3-MPA (n = 10), FK506 (n = 6) or VIVIT (n = 6). \**P*<0.05, \*\**P*<0.01; ns, not significant.

**Figure 4-Figure Supplement 1.**
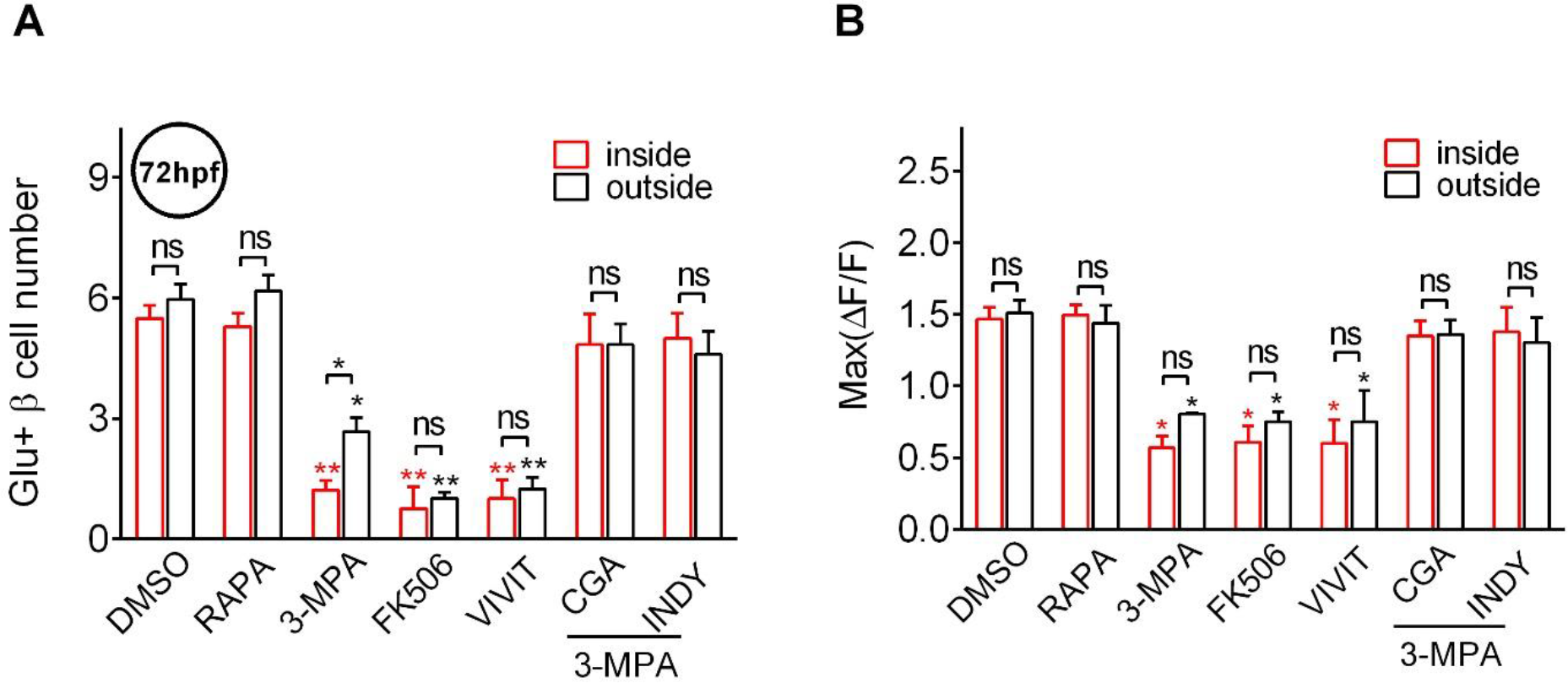
Glucose activates calcineurin/NFAT to finish the final maturation of pancreatic β-cells during the late hatching period. *Tg(ins:Rcamp1.07*) embryos at 60 hpf were treated with different reagents for 9 h and then washed in the normal E3 medium for 3 h before imaging. (A-B) Quantification of glucose-responsive β-cells (*A*) and their maximal ΔF/F in glucose-responsive β-cells (*B*) in the mantle and core of the islet in DMSO-treated controls, rapamycin-treated embryos, 3-MPA-treated embryos, FK506-treated embryos, VIVIT-treated embryos, 3-MPA and CGA co-treated embryos and 3-MPA and ProINDY co-treated embryos at 72 hpf. n = 5-9 embryos for each condition. \**P*<0.05, \*\**P*<0.01; ns, not significant.

**Supplemental Table 1.**
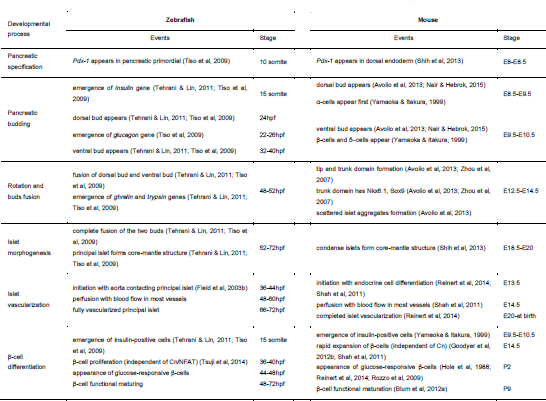
Comparison of the critical events and the time windows in zebrafish and mouse pancreas development.

---

**Supplemental Movie 1. Time-lapse images of glucose-stimulated calcium transients in pancreatic β-cells in a 72 hpf *Tg(ins:Rcamp1.07*) embryo under a wide field microscope.**

**Supplemental Movie 2. Scheme of the dual-colour 2P3A-DSLM and comparison among 1P-SPIM, TPM and 2P3A-DSLM in islet 3D imaging of live zebrafish.**

**Supplemental Movie 3. Representative time-lapse volumetric reconstruction of glucose-stimulated calcium transients in all pancreatic β-cells from 72 hpf *Tg(ins:Rcamp1.07*) embryos observed under the 2P3A-DSLM.**

**Supplemental Movie 4. Volumetric reconstruction of pancreatic β-cells and their nearby blood vessels in living *Tg(ins:EGFP);Tg(flk1:mCherry*) embryos from 36 hpf to 72 hpf.**

**Supplemental Movie 5. Representative time-lapse volumetric reconstruction of glucose-stimulated calcium transients in all pancreatic β-cells from 72 hpf *Tg(ins:Rcamp1.07*) embryos that had been treated with 3-MPA.**

**Supplemental Movie 6. Representative time-lapse volumetric reconstruction of glucose-stimulated calcium transients in all pancreatic β-cells from 72 hpf *Tg(ins:Rcamp1.07*) embryos that had been cotreated with 3-MPA and CGA.**

